# Two-speed genome expansion drives the evolution of pathogenicity in animal fungal pathogens

**DOI:** 10.1101/2021.11.03.467166

**Authors:** Theresa Wacker, Nicolas Helmstetter, Duncan Wilson, Matthew C. Fisher, David J. Studholme, Rhys A. Farrer

## Abstract

The origins of virulence in amphibian-infecting chytrids *Batrachochytrium dendrobatidis* (*Bd*) and *Batrachochytrium salamandrivorans* (*Bsal)* are largely unknown. Here, we use deep nanopore sequencing of *Bsal* and comparative genomics against 21 high-quality genome assemblies that span the fungal Chytridiomycota. *Bsal* has the most repeat-rich genome, comprising 40.9% repetitive elements, which has expanded to more than 3X the length of its conspecific *Bd*. M36 metalloprotease virulence factors are highly expanded in *Bsal* and 53% of the 177 unique genes are flanked by transposable elements, suggesting repeat-driven expansion. The largest M36 sub-family are mostly (84%) flanked upstream by a novel LINE element, a repeat superfamily implicated with gene copy number variations. We find that *Bsal* has a highly compartmentalized genome architecture, with virulence factors enriched in gene-sparse/repeat-rich compartments, while core conserved genes occur in gene-rich/repeat-poor compartments. This is a hallmark of two-speed genome evolution. Furthermore, genes with signatures of positive selection in *Bd* are enriched in repeat-rich regions, suggesting they are a cradle for chytrid pathogenicity evolution, and *Bd* also has a two-speed genome. This is the first evidence of two-speed genomes in any animal pathogen, and sheds new light on the evolution of fungal pathogens of vertebrates driving global declines and extinctions.

## Introduction

*Batrachochytrium salamandrivorans* (*Bsal*) threatens amphibians globally and is currently expanding its geographic range across Europe. It infects highly susceptible fire salamanders, with outbreaks reported in the wild in Germany, Belgium, the Netherlands and Spain (Beukema et al., 2021; Martel et al., 2014). This ecologically important fungal pathogen belongs to the Rhizophydiales order of the Chytridiomycota which includes genera with saprobic free-living as well as pathogenic life histories. For example, *Entophylctis helioformis* and *Homolaphlyctis polyrhiza* are the two closest known relatives to *Batrachochytrium*. However, unlike those amphibian pathogens, *E. helioformis* and *H. polyrhiza* are saprotrophs found on algae and leaf litter and are unable to grow on amphibian skin (Joneson et al., 2011; Longcore et al., 2012).

*Bsal* likely diverged from *Bd* between 30 and 115 million years ago in the Late Cretaceous or early Paleogene, and both species have likely been endemic to Asian salamanders and newts (Urodela) for millions of years. Both species have expanded their ranges in recent time with *Bd* becoming globally established in the early to mid-20^th^ Century while *Bsal* emerged in the Netherlands only in 2010 and spread to naïve European populations (Martel et al., 2014). Since diverging, *Bd* and *Bsal* have evolved to infect different amphibian species and display different pathologies. While *Bd* is a generalist pathogen that infects all three orders of amphibian, *Bsal* has evolved as a specialist pathogen of the Urodela order (newts and salamanders) (Martel et al., 2013), yet is able to survive asymptomatically on amphibians of other orders, potentially contributing to its spread (More et al., 2018). While *Bd* causes hyperplasia (proliferation of cells) and hyperkeratosis (thickening of the stratum corneum), *Bsal* causes multifocal superficial erosions and deep ulcerations in the skin of its host (Martel et al., 2013). The evolutionary route to pathogenicity and the genetic mechanisms underlying host-specificity and pathology in the *Batrachochytrium* genus remain largely unknown.

Evolution shapes genomes unevenly, resulting in both conserved and faster evolving genomic regions. In extreme cases, a phenomenon termed the ‘two-speed genome’ has been identified, whereby rapidly-evolving genes comprise a substantial portion of the genome and is associated with an enrichment of repeat families that are likely contributing to or driving gene variation (Dong et al., 2015; Faino et al., 2016b; Frantzeskakis et al., 2019; Gijzen, 2009; Haas et al., 2009; Lamour et al., 2012; Raffaele et al., 2010; Raffaele & Kamoun, 2012a; Tyler et al., 2006). In plant pathogens with two-speed genomes, fast-evolving regions are enriched for genes that are upregulated *in planta* (Raffaele et al., 2010), have signatures of positive selection (Sánchez-Vallet et al., 2018), and have undergone increased gene family expansions (Raffaele et al., 2010). To date, two-speed genome compartmentalization has been identified in a range of fungal and oomycete plant pathogens (Faino et al., 2016b; Plissonneau et al., 2016; Raffaele et al., 2010; Torres et al., 2020; Q. Wang et al., 2017; Winter et al., 2018). Among the chytrids, *Synchytrium endobioticum* responsible for potato wart disease has been noted to have effector genes within repeat-rich regions (van de Vossenberg et al., 2019). However, there has been no comprehensive analysis or identification of two-speed genomes among the Chytridiomycota to date, or indeed among any animal pathogens.

In 2017, we sequenced *Bsal*’s genome for the first time (R. A. Farrer et al., 2017), discovering it had an expanded genome relative to its closest relatives. We found that *Bsal* has undergone several large protein family expansions, including the M36 metalloproteases that are thought to be involved in the breakdown of amphibian skin and extracellular matrix (Joneson et al., 2011). The M36 metalloprotease family expansions were noted to coincide with an increase in repeat-rich regions; however, that study found only 17% of the genome assembly to be repetitive (R. A. Farrer et al., 2017). We also found evidence that unlike *Bd, Bsal* does not illicit a clear immune response during infection in a shared host species (R. A. Farrer et al., 2017). However, the use of exclusively short-read sequencing limited us to a highly fragmented genome from which we were unable to fully explore its modes of genome evolution and resolve repeat-rich regions. Furthermore, the genomes of only four chytrid species were compared as opposed to the 22 chytrids investigated in the present study. Here, we use long-read nanopore sequencing to more fully understand the genome evolution of *Bsal* underpinning its host-range and pathogenicity.

## Results

### The repeat-driven expansion of *Batrachochytrium salamandrivorans*

Deep nanopore sequencing and genome assembly of *Batrachochytrium salamandrivorans* (*Bsal*) revealed that it has undergone a large genome expansion compared with all known and genome-sequenced Rhizophydiales (**Figure 1**). Notably, the genome of *Bsal* is >3X longer than its closest known relative *B. dendrobatidis* (*Bd*). Our updated genome assembly (version 2; v.2) is a substantial improvement on our previous Illumina-based assembly (version 1; v.1). Notably, v.2 has a total length of 73.3 Mb across 165 supercontigs (*N*_*Max*_ 5.6 Mb, *N*_*50*_ 0.9 Mb) compared with v.1 that is 32.6 Mb across 5,358 contigs (*N*_*50*_ 10.5 kb) (**Table S1**). *Bsal*’s updated genome length elevates it to the second-largest in the Chytridiomycota fungal phyla, after *Cladochytrium polystormum* (81.2 Mb) - a species that is mainly associated with aquatic plants (Czeczuga, Mazalska, et al., 2007; Czeczuga, Muszyńska, et al., 2007; Powell et al., 2018). Our updated gene annotation also revealed slightly higher numbers of predicted protein-coding genes (*n* = 10,867 with a combined length of 16.38 Mb) and was more complete (94.1% of BUSCO core conserved fungal genes) compared to the v.1 assembly (BUSCO = 92.2%). Synteny indicated there were no newly evolved or acquired chromosomes in *Bsal*’s genome, although its genome expansion relative to *Bd* was accompanied by an abundance of chromosomal rearrangements (**Figure S1**).

**Figure 1.**
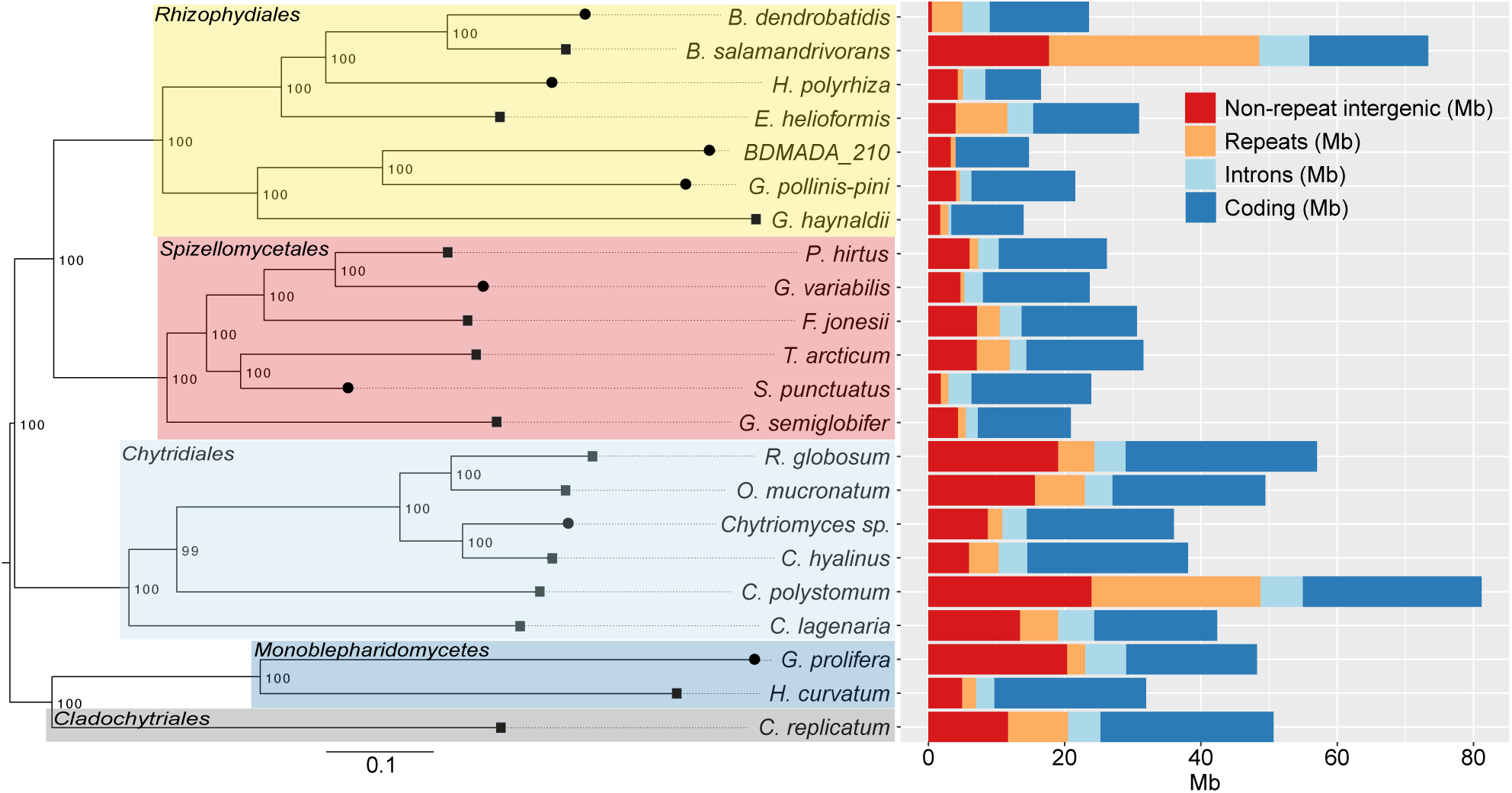
Phylogeny of 22 Chytrids based on multiple alignments of 143 core orthologs (left) next to genome content (right). Tree branches with a round tip have been sequenced using short-read sequencing technologies (Illumina). Tree branches with squares have been sequenced with long-read sequencing technologies (*B. salamandrivorans*: Oxford Nanopore sequencing technology; all others: PacBio sequencing technology). The percentage of 1000 ultrafast bootstrap resampling’s that support the major topological elements in neighbour joining is indicated. The scale bar indicates the number of substitutions per site.

*Bsal* has the most repeat-rich genome of any chytrid sequenced to date, with 40.9% (30 Mb) of the genome predicted to be repetitive (**Figure S2**). *Bsal* has undergone a unique repeat-driven expansion compared with other chytrids including *Bd*, resulting in distinct repeat family profile (**Figure 2**). Repeat content across the Chytridiomycota (as a percentage of genome length) has a positive monotonic correlation with genome length (Spearman’s r_s_ = 0.62, *p* = 0.0019), as do transposable elements (Spearman’s r_s_ = 0.56, *p* = 0.0059). Conversely, repeat content does not correlate with assembly contiguity (N_50_) (Spearman’s r_s_ = 0.19, *p* = 0.39) or degree of fragmentation (number of contigs) (Spearman’s r_s_ = -0.022, *p* = 0.92) (**Figure S3**). Genome length in the Chytridiomycota is therefore a good predictor of repeat-richness.

**Figure 2.**
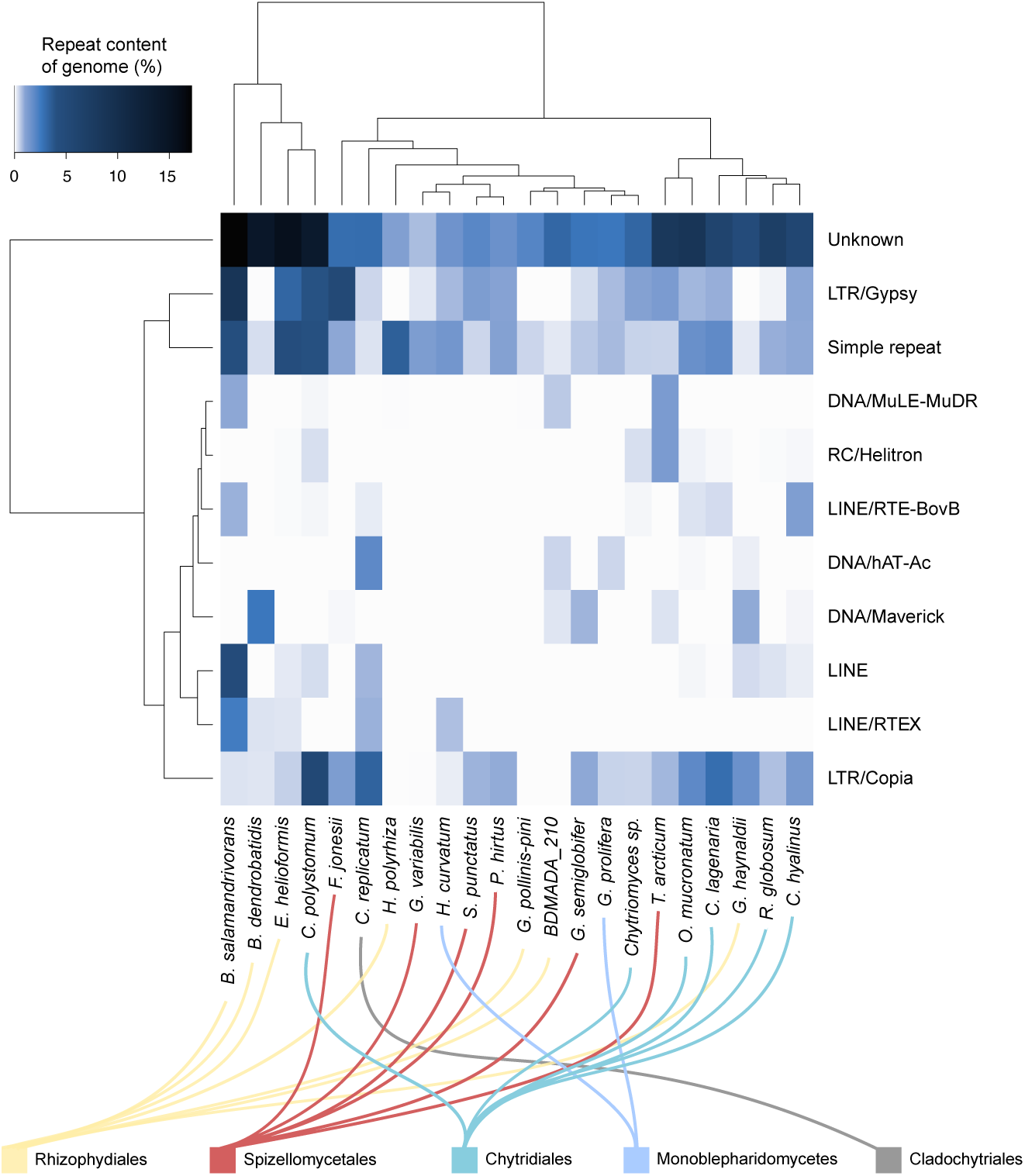
Repeat superfamily abundance in 23 Chytrids. The dendrogram is based on euclidean distances and hierarchical clustering. Only repeat families that have more than 1% abundance in at least one chytrid are included.

Transposable elements (TE), long terminal repeat (LTR) and long interspersed nuclear elements (LINE) retrotransposons are uniquely expanded in the *Bsal* genome compared with *Bd* (**Figure 2**). *Bsal* has the highest overall content of TEs of any chytrid (**Figure S4**). TEs comprise 19.36% of all *Bsal* repetitive content and are not uniformly distributed in the genome but appear in clusters. Conversely, repeats in general, including simple repeats are uniformly distributed (**Figure S5**). *C. polystomum* (the largest genome) has the second highest proportion of TEs (12.3%). All other chytrids have <10% TEs (geometric mean: 3.43%, *σ*: 2.52%; excluding *Bsal* and *C. polystomum*), indicating that TE content associates with genome expansions. LTRs are the second most abundant repeat family in *Bsal* (6.6 Mb; 9% of genome), most of which (97%) are Gypsy elements. Gypsy repeats are far less common in other Rhizophydiales including *Bd* (4.8 kb; 0.02% of genome), *H. polyrhiza* (absent), and *E. helioformis* (988 kb; 3.2%). Similarly, LINEs make up 6.4 Mb (8.8%) of the *Bsal* genome (**Table S2**), yet are not detected in most of the other genomes belonging to the Rhizophydiales including *H. polyrhiza*, BdMADA_210 (amphibian-associated chytrid recovered from Madagascar) and *Globomyces pollinis-pini* (a saprotrophic chytrid found in aquatic habitats (Pm et al., 2008)). The three remaining Rhizophydiales species (*Bd, E. helioformis* and *G. haynaldii*) have only low numbers of LINE elements (0.28%, 0.6% and 0.47% of genome, respectively).

### The ‘two-speed’ compartmentalized genome of *Batrachochytrium salamandrivorans*

Analysis of chytrid intergenic distances revealed *Bsal* has a compartmentalized, bipartite genome. Disparate flanking intergenic region (FIR) lengths for several gene families and biological functions were identified across the Chytridiomycota (**Figure 3, Figure S6**). To assess differences between FIR lengths and gene categories, we characterised four groups or quadrants partitioned by the 5’ and 3’ median intergenic distances (upper-left; Q_UL_, upper-right; Q_UR_, lower-left; Q_LL_ and lower-right; Q_LR_) and tested for enrichment of genes using hypergeometric tests (HgT) and χ^2^ tests. The set of all *Bsal* genes were evenly dispersed across quadrants (Q_UL_ = 2,269 genes, Q_UR_ = 2,998 genes, Q_LR_ = 2,268 genes, and Q_LL_ = 2,998 genes). However, the subset of core conserved genes (CCGs) is enriched in Q_LL_ (HgT *p* = 5.88E-6, χ^2^ test *p* = 7.16E-6). Furthermore, M36 protease genes, other genes encoding proteins with secretion signals and genes encoding small secreted proteins (SSPs) were all enriched in Q_UR_ according to HgT (*p* = 1.2E-38, *p* = 5.42E-92, *p* = 9.13E-7, respectively) and χ^2^ tests (*p* = 1.93E-43, *p* = 5.39E-102, *p* = 4.4E-7, respectively). This is a hallmark of a two-speed genome (**Table S3B-C)**). Accordingly, *Bsal* has 3.5X more repetitive sequence and 4.5X more TE in Q_UR_ compared with Q_LL_ based on gene identities in 10 kb non-overlapping windows (**Table S3K**).

**Figure 3.**
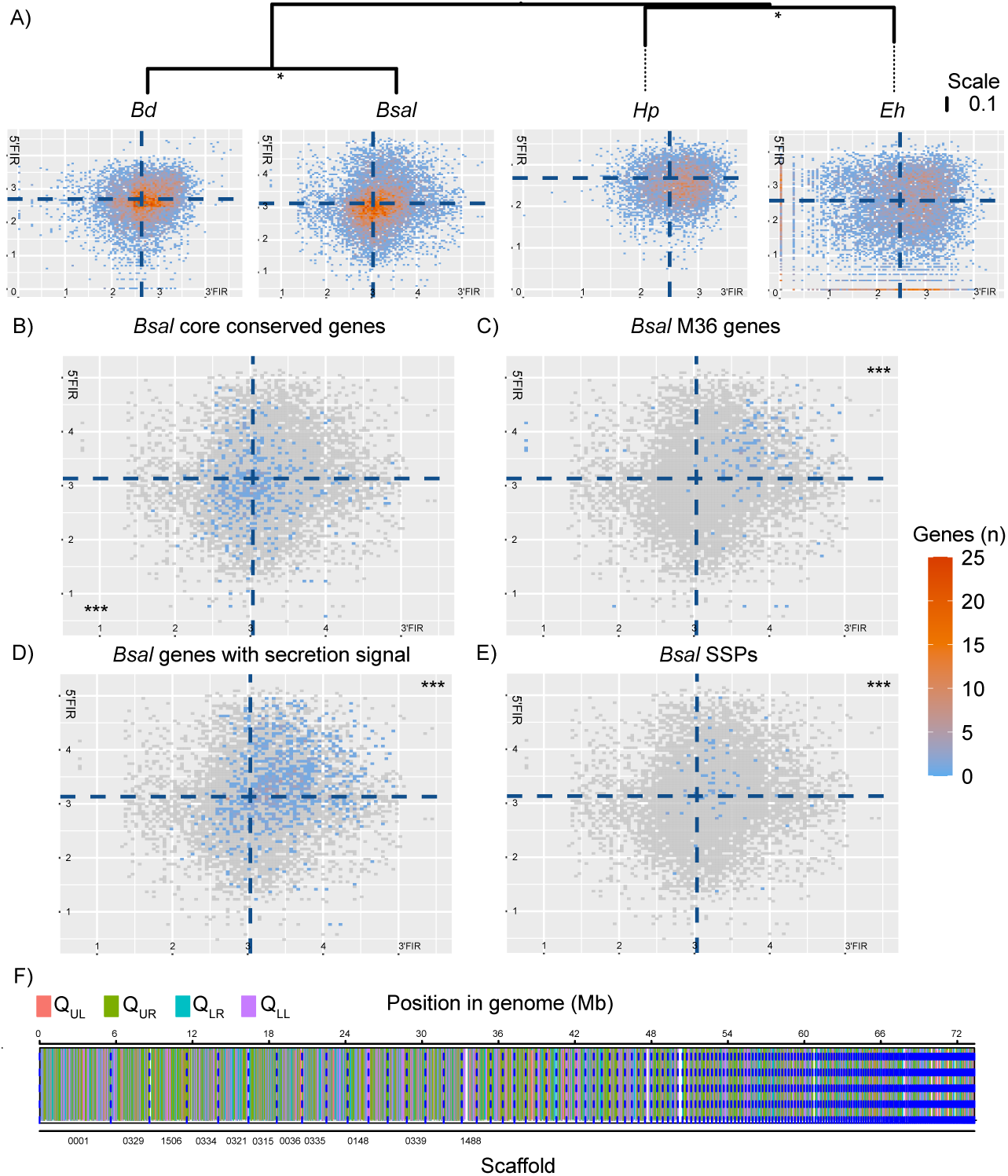
The two-speed genome of *Bsal*. **A)** a phylogenetic tree of *Bsal* and its three closest relatives: *Bd, Hp*, and *Eh* constructed using a core-ortholog multiple alignment and RAxML. Vertical branch lengths indicating the mean number of nucleotide substitutions per site. Asterisks indicate 100% bootstrap support from 1,000 replicates. Density plots of intergenic distance for all non-terminal protein coding genes are shown (measured as log_10_ length of 5’ and 3’ flanking intergenic distances; FIRs), with the median values shown in dotted blue line. **B-E)** density plots of intergenic distance for *Bsal* gene categories. Individual plots are non-terminal **B)** core-conserved genes, **C)** M36 metalloprotease, **D)** secreted genes and **E)** small secreted genes respectively. The median intergenic distance for all genes is shown as a dotted blue line. Asterisks indicate enrichment of genes in one of the four quadrants based on the median intergenic distances (hypergeometric test, α-level = 0.01). **F)** shows the positions of 10kb windows assigned to the quadrants in the genome.

FIR lengths are significantly longer for M36 proteases, genes encoding proteins with secretion signals and SSPs compared to overall mean intergenic distances in *Bsal* (Wilcoxon rank-sum tests: *p* = 1.42E-61, 2.45E-121 and 2.50E-07, respectively) (**Table S3D**). Mean intergenic distances for M36 proteases, genes encoding proteins with secretion signals and SSPs are also significantly longer than CCGs (Wilcoxon rank-sum tests: *p* = 1.46E-61, 8.54E-72 and 2.4E-13, respectively). Separately, CCGs are flanked by significantly shorter intergenic regions than the genome-wide average (Wilcoxon rank-sum test *p* = 2.03E-09) (**Figure 4, Table S3D)**. Intriguingly, most chytrids (18 out of 23) had an enrichment of CCGs in Q_LL_ and nearly half (10 out of 23) had an enrichment for genes encoding proteins with secretion signals in Q_UR_ (HgT and χ^2^ test *q* < 0.01), indicating those are common features of Chytidiomycota evolution. However, *Bsal* has the most significant enrichment of genes with secretion signals in Q_UR_ of any chytrid tested (HgT *p* = 5.42E-92, χ^2^-test *p* = 5.39E-102) (**Table S3B-C**), while *Bd* had the second strongest enrichment (HgT *p* = 9.75E-67, χ^2^-test *p* = 4.18E-74).

**Figure 4.**
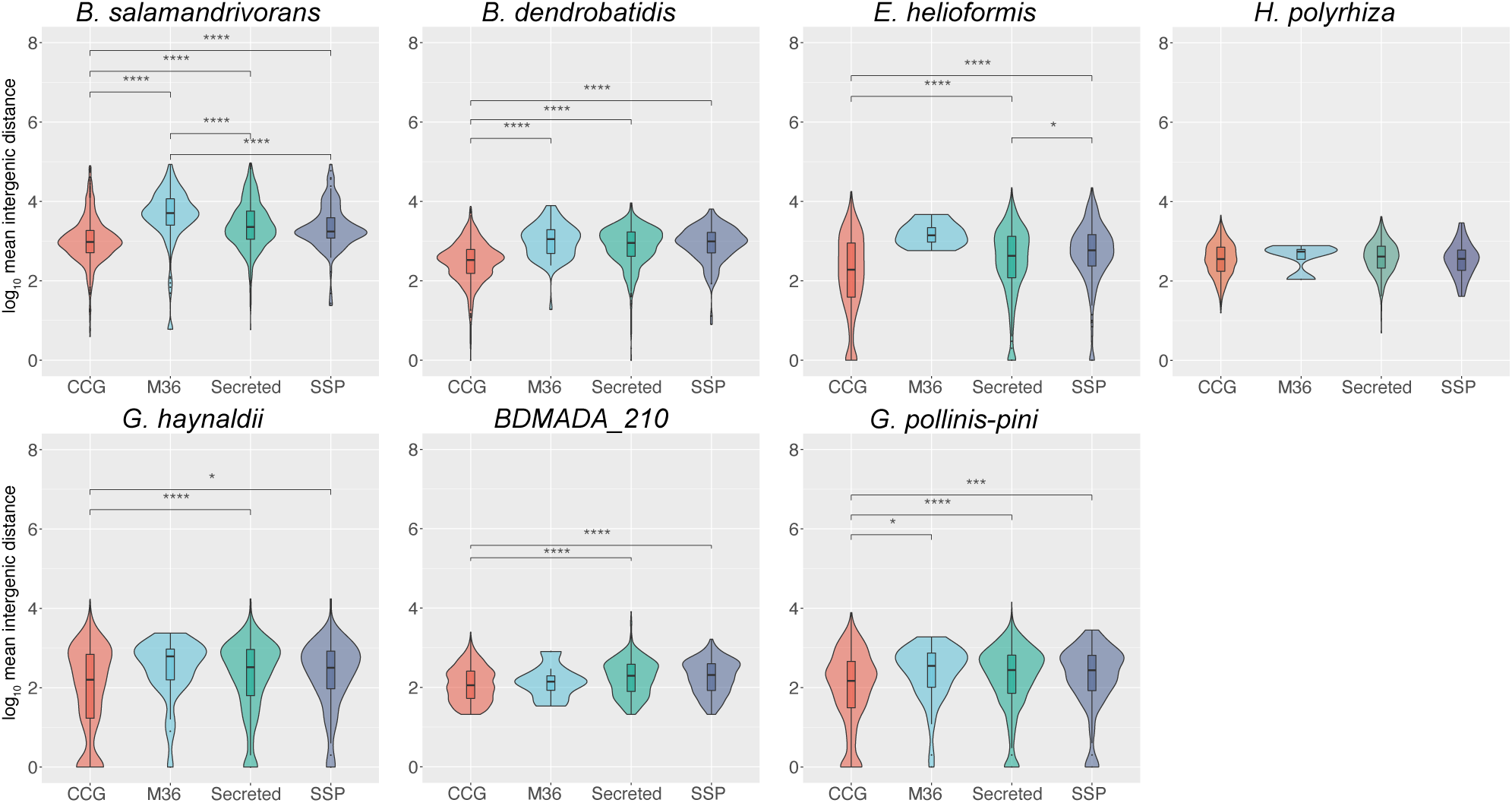
Rhizophydiales Intergenic Distance Analysis. Log_10_ mean intergenic distances of core-conserved genes (CCGs), SSPs (small secreted proteins; <300 aa and ≥ 4 cysteines), secreted proteins and M36 metalloproteases for chytrids of the clade Rhizophydiales. Boxplots indicate median and interquartile range. To determine statistically significant differences of the log_10_ mean intergenic distances for pairs of different features of interest, a Wilcoxon rank-sum test was performed. P-values were Bonferroni adjusted with an α-level of 1 and n = 6. *: p ≤ 0.0017, **: p ≤ 0.00017, ***: p ≤ 1.7E-5, ****: p ≤ 1.7E-6

FIR based quadrants are present throughout *Bsal*’s genome and not exclusively from individual chromosomes or large sub-chromosomal or subtelomeric regions (**Figure 3F**). Similarly, M36 metalloproteases are encoded throughout the *Bsal* and *Bd* genomes (**Figure S1**). Of the 28 contigs that feature a (TTAGGG)_n_ terminal telomeric repeat, 14 feature clusters of up to 10 M36s or gene with secretion signal in those subtelomeric regions. Six contigs overall deviate from the null hypothesis of 25% of genes populating each quadrant (χ^2^-test for goodness of fit; **Table S4**), three of which (scaffolds 94, 329 and 334) are >1 Mb in length. Twelve contigs were enriched for one of the quadrants (HgT *p* < 0.01) including Q_UR_ (*n* = 9) and Q_LL_ (*n* = 3) and no enrichments were found for either Q_UL_ or Q_LR_ (**Table S4**). The longest stretch of consecutive Q_UR_ genes is 16 (128 kb), Q_LL_ genes is 13 (33 kb), Q_LR_ genes is 5 (25 kb), and for Q_UL_ it is only 4 (16 kb; **Table S5A**).

The probabilities for each set of consecutive genes from any given quadrant was calculated using a custom discrete-time pattern Markov chain approach, which identified 35 contigs with significant consecutive gene counts (*p* < 0.01) from either Q_UR_ (*n* = 15) or Q_LL_ (*n* = 20; **Table S5B**). The two most significant of these were scaffold0320 with 16 consecutive genes in Q_UR_ (total genes on contig = 122, *p* = 1.44E-07) and scaffold0334 with 15 consecutive genes in Q_UR_ (total genes on contig = 380, *p* = 1.71E-06). The 16 consecutive Q_UR_ genes on contig320 included 6 genes with secretion signals (three belonged to Tribe 31 (unknown function), and two belonged to Tribe 17 with a S1-P1 nuclease domain (PF02265.16)). The 15 consecutive Q_UR_ genes on scaffold0334 included only 1 gene with a secretion signal (Tribe 536, with a kelch4 galactose oxidase central domain (PF13418.6)).

Genes with long intergenic distances are associated with positive selection in *Bd*. While all reported isolates of *Bsal* to date are clonal, there have been substantial sampling efforts for *Bd* revealing five genetically diverse lineages, providing an opportunity to explore intra-population genetic variation and associations with intergenic distances, which is an opportunity not currently available for *Bsal*. By calculating *dN*/*dS* (ω) for every *Bd* gene in each lineage, we discovered that genes with a signature of positive selection (ω > 1) were significantly enriched in Q_UR_ for each lineage (HgT *p* < 2.59E-10, χ^2^-test *p* < 6.5E-11) (**Table S3E)**. Notably, genes with ω > 1 and secretion signals were enriched in Q_UR_ for each *Bd* lineage (HgT *p* < 8.48E-13, χ^2^-test *p* < 3.64E-14). This is consistent with both Batrachochytrids (*Bd* and *Bsal*) having two-speed genomes.

Recently, it was shown that ricin-like B lectins play a crucial role in the initial stages of pathogenesis in *Bsal* (Y. Wang et al., 2021). Previous studies have also found that these lectins are expressed during exposure of *Bsal* to salamander skin (R. A. Farrer et al., 2017). Of the two ricin-like B lectins identified in this study, one of them, BSLG_002240, can be found in Q_UR_, providing evidence that virulence genes are indeed sequestered to this dynamic compartment.

*Bsal* encodes the greatest number of secreted proteins (*n* = 1,047) in the Rhizophydiales. Clustering secreted proteins by amino acid sequence for *Bsal* and its three closest relatives *Bd, Hp* and *Eh*), revealed 854 distinct secreted tribes, including the M36 metalloproteases that is the largest (*n* = 167, of which 142 belong to *Bsal*; Tribe 1). The ten largest secreted tribes encompass nearly a quarter of all secreted proteins of *Bsal, Bd, Hp* and *Eh* (24.08%; *n* = 593) (**Table S3J)**. Notable gene tribes included the M36s (Tribe 1), polysaccharide deacetylases (Tribe 4), tyrosinases (Tribe 6), aspartyl proteases (Tribe 7), phosphate-induced proteins (Tribe 8) and lipases (Tribe 10), each of which may be involved in pathogenicity. *Bsal* genes in Tribes 1, 4, 8 and 10 are enriched in Q_UR_. *Bd* genes in Tribes 1, 2, 3, 5, 7 and 9 are enriched in Q_UR_ (**Figure 5, Table S3I-J, Table S6)**.

**Figure 5.**
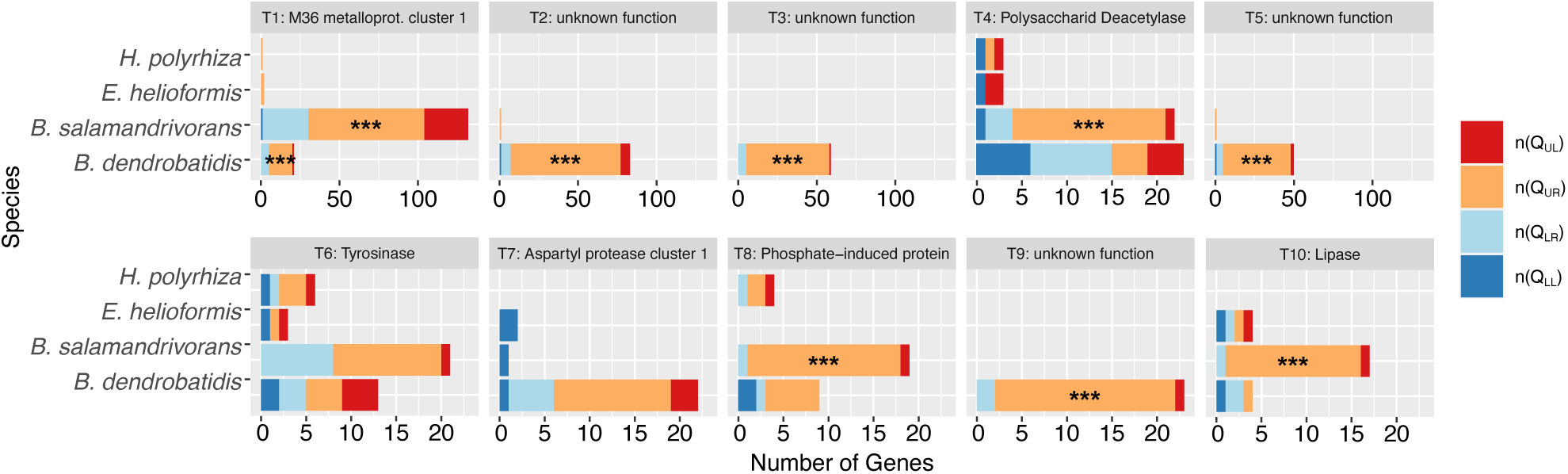
Secreted protein tribes in *Bsal, Bd, Eh* and *Hp*. The 10 biggest secreted protein tribes identified by clustering all predicted secreted genes using MCL found in one or more of the 4 chytrids *Bsal, Bd, Eh*, and *Hp*. Significance of enrichment was determined using hypergeometric tests. Q_UR_ = gene-sparse region, Q_LL_ = gene-rich region. Asterisks indicate enrichment of genes in one of the four quadrants (hypergeometric test, p_adjusted_ < 0.00063).

### M36 metalloprotease expansion linked to transposable elements

*Bsal* encodes the most M36 metalloproteases (*n* = 177) of any other chytrid (**Fig. S1**), which is >5X more than *Bd* encodes (*n* = 35) (Joneson et al., 2011). M36 metalloproteases in *Bsal* can be divided into six species-specific families (*Bsal* M36 family 1-6) and two more evolutionary conserved families based on sequence similarity and a gene tree (**Figure 6**). The largest M36 sub-family (*n* = 70; *Bsal* M36 family 6) was previously named *Bsal* G2M36 (R. A. Farrer et al., 2017) and is uniquely associated with two repeat families including the transposable LINE element rnd2 family2 (**Table S2, Table S7**). The majority of *Bsal* M36 family 6 genes are flanked by this LINE element upstream (*n* = 59/70; 84.3%; *p* = 3.33E-99) and flanked by an uncharacterised repeat rnd1 family182 downstream (53/70; 75.7%; *p* = 8.56E-75), as well as 67.1% of *Bsal* M36 family 6 are flanked either side by both repeats. No other *Bsal* M36 family has a flanking rnd1 family182 and only 2 *Bsal* M36s have a flanking rnd2 family2 (M36 families 1 and 3). The 8.8% of *Bsal*’s genome comprising LINE elements is therefore associated with its genome expansion and gene family expansion of putative virulence factors.

**Figure 6.**
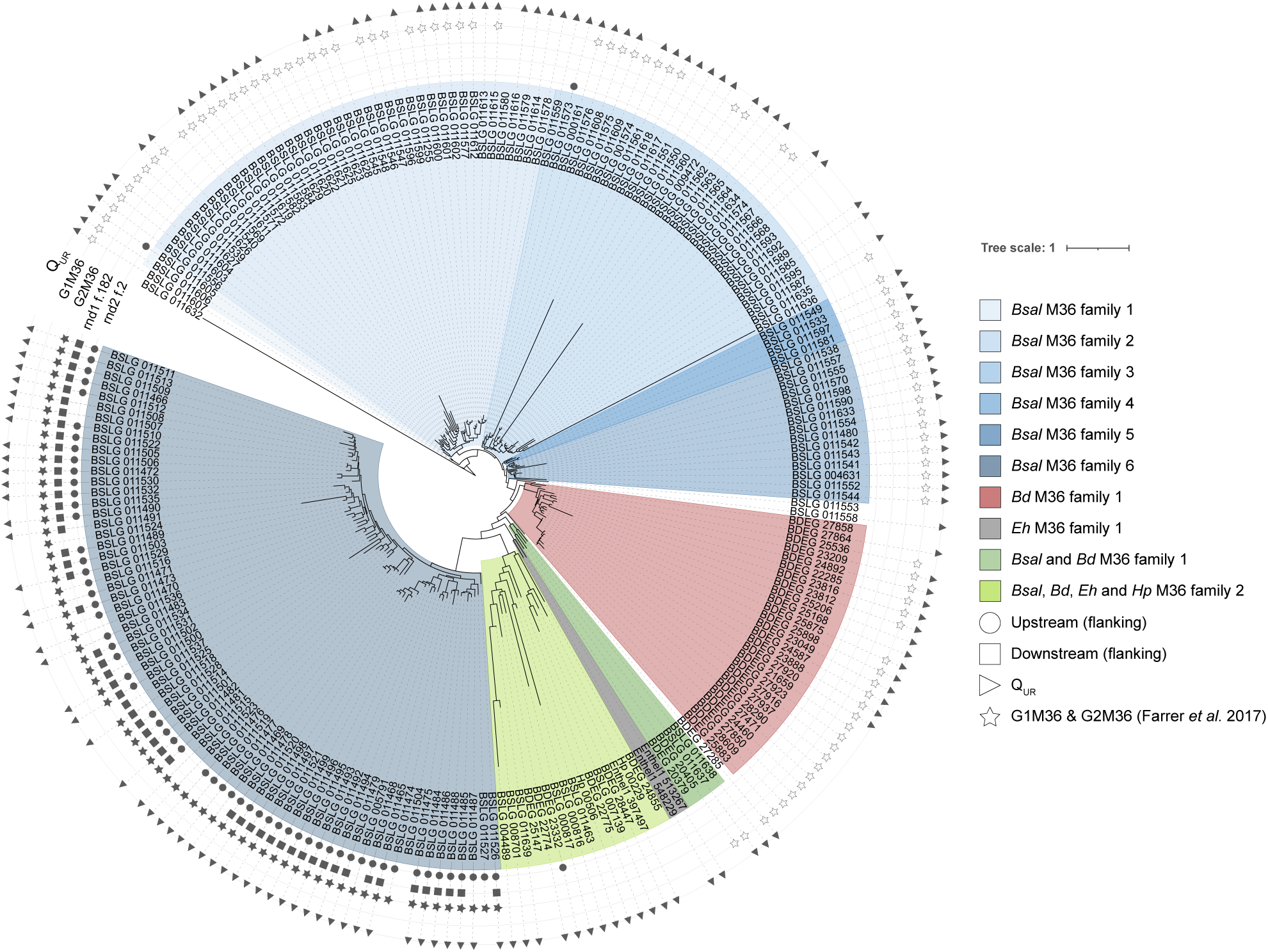
Phylogenetic tree inferred using RaxML from protein alignments of all identified M36 proteins in four chytrids. RaxML-inferred tree of all known M36 metalloproteases in Bsal, Bd, Eh and Hp. The branch lengths (Tree scale) indicate the mean number of nucleotide substitutions per site. Upstream and downstream flanking repeat families, rnd2 family2 (LINE) and rnd1 family182 (Unknown) are indicated as circles and squares, respectively. Localization of the M36 metalloprotease genes in the gene-sparse region is indicated by triangles. G1M36 and G2M36 (as previously defined (R. A. Farrer et al., 2017)) are marked by star outlines and filled star shapes, respectively. Gene IDs: Enthel1 = *E. helioformis, Hp* = *H. polyrhiza*, BSLG = *B. salamandrivorans* and BDEG = *B. dendrobatidis*.

*Bsal* M36-associated repeats are highly enriched upstream and downstream of the M36 metalloprotease coding genes (rnd2 family2 upstream of M36s: p = 8.56E-75; rnd1 family182 downstream of M36s: 3.33E-99) and genes with secretion signals (rnd2 family2 upstream of secreted: *p* = 4.50E-28; rnd1 family182 downstream of secreted: p=1.48E-42; **Table S3F-H**). Rnd1 family182 and rnd2 family2 are disproportionately found in gene-sparse/repeat-rich compartments of the genome (Q_UR_: p_rnd2 family2_ = 2.53E-25, p_rnd1 family182_ = 1.16E-11). Rnd2 family 2 has a homologous repeat family in *Bd* (rnd1 family109), which is also classified as a LINE. However, this repeat family is only present upstream of 1 M36 metalloproteases and 8 genes with secretion signals in *Bd*. Rnd1 family182 has no homologous repeat family in *Bd*. We found only 66/734 *Bsal* repeat families had homologs in *Bd* suggesting that *Bsal*’s two-speed genome is largely driven by repeat families that have emerged since it speciated with *Bd*.

## Discussion

The chytridiomycosis panzootic has been identified as one of the key drivers of global amphibian declines, contributing to earth’s sixth mass extinction. Since the discovery of *Bd* (Berger et al., 1998) and more recently *Bsal* (Martel et al., 2013), efforts have been made to understand their evolution and mechanisms of pathogenicity and virulence. *Bsal*’s virulence is likely shaped by an “arms race” between host and pathogen, resulting in large and diverse families of proteolytic enzymes for skin and extracellular matrix destruction (R. A. Farrer et al., 2017; Fisher et al., 2021; Papkou et al., 2019). Here, we assemble and annotate an improved *Bsal* genome assembly and perform comparative genomics across the Chytridiomycota, discovering that both *Bsal* and *Bd* have hallmarks of two-speed genomes.

*Bsal* has an extremely repeat-rich genome (40.9%) compared with most fungal species, which typically range from ∼5-35% (Wöstemeyer & Kreibich, 2002). However, there are other fungal pathogens with even greater repeat content (>62%) such as *Venturia* and *Blumeria* species (Cam et al., 2019; Castanera et al., 2016; Frantzeskakis et al., 2018; Peter et al., 2016; Wöstemeyer & Kreibich, 2002). Many of *Bsal’*s repeats are transposable elements (TEs) (19.36%) which underpin the genome expansion described in other species including the fungal wheat pathogen *Zymoseptoria tritici*, the barley powdery mildew, *Blumeria graminis* f.sp. *hordei*, the oomycete causative agent of potato blight, *Phytophthora infestans* and the symbiotic fungus *Cenococcum geophilum* (Oggenfuss et al., 2021; Peter et al., 2016; Raffaele & Kamoun, 2012a; Spanu et al., 2010). TEs are abundant in the genomes of various fungal pathogens, comprising 36% of the genome for plant pathogen *Leptosphaeria maculans*, 64% of the genome for *B. graminis* and 74% for *P. infestans* (Grandaubert et al., 2014; Raffaele & Kamoun, 2012a; Spanu et al., 2010). TEs in *Bsal*’s closest known relative *Bd*, however, only comprises 3.37% of its genome, suggesting they have expanded recently in *Bsal* and that the differences in genome size in the batrachochytrids associates with TE expansion in *Bsal*. The most abundant TE family in *Bsal* is LTR/Gypsy, which is almost absent in *Bd* (0.02%). LTR/Gypsy has been previously identified as a driver of genome expansion and also has been implicated with adaptation along environmental gradients and under stress conditions (Marcon et al., 2015; Pietzenuk et al., 2016; Y. Wang et al., 2018; Wos et al., 2021; Zhang et al., 2020). More broadly, we found a strong correlation between TE and repeat content with genome size across the Chytridiomycota. To our knowledge, this is the first time that a statistical correlation between TE and repeat content with genome size has been shown for an order of fungi, with previous studies finding correlations in insects, chordates, larvaceans and tetrapodes, but not fungi or fungal orders specifically (Canapa et al., 2015; Kidwell, 2002; Naville et al., 2019).

Host-pathogen interactions exerts strong selective pressures, leading to adaptive co-evolution (Papkou et al., 2019; Tellier et al., 2014). Under Red Queen dynamics, both host and pathogen constantly adapt to the ongoing selective pressures imposed by their coevolutionary interactions, with the host’s immune system evolving to detect and mount defences against the pathogen and the pathogen evolving to colonize the host (Brockhurst et al., 2014; Cook et al., 2015; Papkou et al., 2019; Torres et al., 2020; Van Valen, 1973). Pathogens with fluid genotypes may co-evolve more effectively with their host and adapt to new hosts more ably, thereby outcompeting other lineages with less plastic genomes (“clade selection”) (Dong et al., 2015; Raffaele & Kamoun, 2012b; Torres et al., 2020). Many filamentous fungal plant pathogens and oomycetes have bipartite genome architectures with gene-sparse/repeat-rich compartments enriched with effector genes (such as those coding for secreted proteins that function outside of the organism they were synthesised in), acting as “cradles of adaptive evolution” (Dong et al., 2015; Frantzeskakis et al., 2019; Haas et al., 2009; Raffaele et al., 2010; Raffaele & Kamoun, 2012b). These repeat-rich/gene-sparse compartments are associated with higher evolvability and genome plasticity and are often enriched in TEs, feature structural and copy number variations and are enriched with genes under positive selection (Croll & McDonald, 2012; Faino et al., 2016b; Grandaubert et al., 2014; Haas et al., 2009; Plissonneau et al., 2016; Raffaele et al., 2010; Rouxel et al., 2011; Schrader & Schmitz, 2019). Conversely, gene-rich/repeat-sparse compartments are enriched in core conserved genes (Raffaele et al., 2010). This two-speed genome therefore provides an evolutionary solution for high evolvability in some parts and conservation in others, providing genome plasticity while reducing the risk of excessive deleterious mutations in essential genes.

Here we show that the batrachochytrids have such a bipartite “two-speed” genome compartmentalization, which is especially pronounced in *Bsal*. The gene-sparse compartment in both batrachochytrids are enriched in putative effector genes encoding secreted proteins, metalloproteases, and ricin B-like lectins, implicated with chemotaxis, adhesion, and the early stages of a pathogenesis (R. A. Farrer et al., 2017; Gao et al., 2019; Jousson et al., 2004; Ramakrishna Rao & Shanmugasundaram, 1970; Shende et al., 2018; Y. Wang et al., 2021; Xu et al., 2020). In *Bd*, gene-sparse compartments are enriched for genes with signatures of positive selection, indicating that the gene-sparse compartment is a hotspot of adaptive evolutionary processes in batrachochytrids.

M36 metalloproteases are thought to break down the amphibian skin and extracellular matrix during infection of the host (R. A. Farrer et al., 2017). We discovered that large gene families of *Bsal* M36 metalloproteases (such as family 6) are enriched for flanking LINE elements. Furthermore, LINEs are recognized as a source of gene duplications and implicated with genetic novelty where duplicated genes can evolve new functions (Chen et al., 2013; Wicker et al., 2007). Of all the chytrid genomes analysed, only *Bsal* and *Bd* have groups of TEs enriched around putative virulence genes, a further hallmark of two-speed genomes.

In fungi, repeats in genomes are targeted for mutation *via* the repeat induced point mutation (RIP) mechanism, which protects the genome from duplications and transposable element proliferation. In *Bd*, RIP is thought to be absent (van de Vossenberg et al., 2019). Absence of RIP is generally associated with a uniform distribution and with the erosion of compartmentalization of TEs, something that has been found in genomes lacking a compartmentalized structure before (so-called one-speed genomes) (Frantzeskakis et al., 2018, 2019). It is unknown if RIP occurs in *Bsal*. If RIP is absent, the observed compartmentalization of TEs, especially in the context of the uniform distribution of repeats overall, might be achieved by other mechanisms of TE silencing, such as histone modification and methylation (Deniz et al., 2019). The strong association of putative effector genes with TEs and their uneven distribution in *Bsal* suggests that TEs are actively and passively shaping its genome architecture, as well as driving higher evolvability of compartments enriched in virulence factors (Bao et al., 2017; Dong et al., 2015; Faino et al., 2016a; Frantzeskakis et al., 2018; Grandaubert et al., 2014; Raffaele & Kamoun, 2012b; Rouxel et al., 2011; Schrader & Schmitz, 2019).

For the first time, we discover that two important pathogens of vertebrates, *Bsal* and *Bd*, have the hallmarks of two-speed genomes. Such bipartite genome architectures found in several plant pathogens is therefore not limited to plant-pathogens. In the batrachochytrids, their two-speed genomes underpins size and content of their genome, with genes likely to be involved in pathogenicity enriched within genomic compartments that allow for their rapid adaptive evolution.

## Figure and Table legends

**Figure S1.** Synteny plots between **A)** *Bsal* and its three closest relative Bd, Hp and Eh, and **B)** *Bsal* v.1 genome assembly and *Bsal* v.2 genome assembly. The positions of M36 metalloproteases are indicated as blue squares. The phylogenetic tree of *Bsal* and its three closest relatives was constructed using core-ortholog multiple alignment and RAxML (with WAG transition matrix). Branch lengths indicating the mean number of nucleotide substitutions per site. Asterisks indicate 100% bootstrap support from 1,000 replicates.

**Figure S2.** Overall repeat content (%) of all 22 chytrids. Overall repeat content in percent of all 22 chytrids excluding lower-scoring matches.

**Figure S3.** Linear regression plots with Spearman correlation coefficients. **A)** Correlation plot of transposable element content (%) and genome length (bp) for 22 chytrids, **B)** repeat content (%) and the genome length, **C)** repeat content (%) and N%) and **D)** repeat content and the number of contigs. R-squared (R^2^), equation of the linear equation and Spearman’s rank correlation coefficient are indicated in each plot. Data points are blue, the linear regression line is red, the confidence interval is grey.

**Figure S4.** Overall content of transposable elements (%) of all 22 chytrids. Overall content of transposable elements in percent of all 22 chytrids excluding lower-scoring matches (stringent criterion).

**Figure S5.** Repeat and TE distributions in the *Bsal* genome.

**Figure S6.** Density plots for all 22 chytrids analysed. Density plots of intergenic distances for all non-terminal protein coding genes are shown (measured as log_10_ length of 5’ and 3’ flanking upstream and downstream intergenic distances), with the median values shown in dotted blue line.

**Table S1.** Whole genome assembly statistics. All genome assemblies apart from V1 Allpaths (R. A. Farrer et al., 2017), are *de novo* long read assemblies generated in this study. V2 Canu default settings Pilon corrected = *Bsal* assembly v2.0 (our chosen assembly, highlighted in blue).

**Table S2.** Repeat superfamily distributions of the 22 chytrids based on RepeatModeller classifications. Lower-scoring matches were excluded. All values are in base pairs (bp).

**Table S3.** Parameters and *p*-values of hypergeometric tests, χ^2^-tests and Wilcoxon rank sum tests. For all enrichment tests, the four quadrants (Q_UL_, Q_UR_, Q_LR_ and Q_LL_) are based on the 5’ and 3’ median log_10_ intergenic distances.

**A)** Number of M36 metalloprotease genes, genes coding for secreted proteins, small secreted protein (SSP) genes and core-conserved protein (CCG) genes in all 22 chytrids. SSPs are defined as either having fewer than 200 amino acids (aa; all chytrids) or proteins with fewer than 200 aa and ≥ 4 cysteines (for Rhizophydiales).

**B)** *p*-values of hypergeometric tests for enrichment of M36 metalloproteases, secreted proteins, core-conserved genes and small secreted proteins (defined as smaller than 200 aa and ≥4 cysteines for Rhizophydiales and only as smaller than 200 aa for the rest) in the four quadrants for all 22 chytrids. Significant *p*-values (p_adjusted_ < 0.00063; α-level = 0.01, *n* = 16) are highlighted in blue.

**C)** *p*-values for χ^2^-tests for enrichment of M36 metalloproteases, secreted proteins, core-conserved genes and small secreted proteins (defined as smaller than 200 aa and ≥4 cysteines for Rhizophydiales and only as smaller than 200 aa for the rest) in the four quadrants for all 22 chytrids. Significant p-values (p_adjusted_ < 0.00063; α-level = 0.01, *n* = 16) are highlighted in blue.

**D)** Wilcoxon rank-sum test results for distributions of all mean upstream and downstream log_10_ intergenic regions compared to mean upstream and downstream log_10_ intergenic regions of M36 metalloproteases, secreted proteins, core-conserved genes and small secreted proteins (defined as smaller than 200 aa and ≥4 cysteines for Rhizophydiales and only as smaller than 200 aa for the rest) in all 22 chytrids. **** = 1e-04, *** = 0.001, ** = 0.01, * = 0.05, ns = 1. Significant *p*-values (p_adjusted_< 3.0303E-05; α-level = 0.01, *n* = 330) are highlighted in blue.

**E)** Enrichment of M36 metalloproteases, secreted proteins, core-conserved genes and small secreted proteins in *Bd* among the four Quadrants was calculated by hypergeometric tests and χ^2^-tests (*p*-values shown). Significant *p*-values (p_adjusted_ < 0.0005; α-level = 0.01, *n* = 20) are highlighted in blue.

**F)**Enrichment of rnd2 family2 (LINE) and rnd1 family182 (unknown) repeat families in the four quadrants in *Bsal* was calculated using hypergeometric tests and for χ^2^-tests (*p*-values shown). Significant *p*-values (p_adjusted_ < 0.0025; α-level = 0.01, *n* = 4) are highlighted in blue.

**G)** Enrichment of repeat-families upstream and downstream of M36 metalloproteases in *Bsal. p*-values are calculated for hypergeometric tests and χ^2^-tests. Repeat families were annotated according to RepeatModeller classifications. Significant *p*-values (p_adjusted_ < 0.000083; α-level = 0.01, *n* = 120) are highlighted in blue.

**H)** Enrichment of repeat-families upstream and downstream of secreted protein coding genes in *Bsal. p*-values are calculated for hypergeometric tests and χ^2^-tests. Significant *p*-values (p_adjusted_< 0.00001639; α-level = 0.01, *n* = 610) are highlighted in blue.

**I)** Enrichment of the 10 largest secreted tribes in the four quadrants was calculated for *Bsal, Bd, Eh* and *Hp. p*-values were determined using hypergeometric tests. Significant *p*-values (p_adjusted_ < 0.00063; α-level = 0.01, *n* = 16 (four quadrants and four species for each tribe)) are highlighted in blue.

**J)** Numbers of genes in each secreted Tribe and the number and names of assigned PFAMs. NA = Non applicable.

**K)** Details of 10 kb non-overlapping windows categorised according to internal gene Quadrants. Windows with no predicted genes or only terminal genes were considered uncharacterised (Q_unchar._).

**Table S4.** Quadrant enrichment and distribution on the chromosomes. The enrichment tests for the four quadrants Q_UL_, Q_UR_, Q_LR_ and Q_LL_ are based on the 5’ and 3’ median log_10_ intergenic distances.

**A)** Enrichment of genes belonging to one of the 4 quadrants on each chromosome was calculated using Hypergeometric tests, testing if there are more genes in a given quadrant on that chromosome than would be expected for the overall number of genes on the chromosome, given the number of genes in the quadrants in the entire genome. The significance level was adjusted to *p* < 0.0025 (α-level = 0.01, *n* = 4). Significant enrichments are highlighted in blue.

**B)** χ^2^-test for goodness of fit of quadrant distribution on chromosomes. Significant deviations from the distribution of numbers of genes in quadrants from the expected 25% each are highlighted in blue. The significance level is Bonferroni adjusted to *p* < 0000633 (α-level = 0.01, *n* = 158). Obs = observed count; Exp = expected count.

**Table S5.** Consecutive gene counts and discrete-time Markov Chain probabilities.

**A)** Consecutive gene counts for the four quadrants (Q_UL_, Q_UR_, Q_LR_ and Q_LL_) are based on the 5’ and 3’ median log_10_ intergenic distances. Start and end position of the block of consecutive genes are indicated.

**B)** Probabilities for the number of consecutive genes found in the four quadrants. Probabilities of finding consecutive genes of length k on contigs with n genes, computed using discrete-time pattern Markov chains. Significant (Sig) *p*-values (p<0.01) are marked with *.

**C)** Longest number of consecutive genes in each of the four quadrants for each chytrid.

**Table S6.**Matching tribes of *Bsal* assemblies v2.0 and v1.0. Gene IDs of genes assigned to the respective tribes are denoted in *Bsal* v2.0 Gene ID and *Bsal* v1.0 Gene ID. Note that IDs in the same row are not homologous.

**Table S7.**M36 Metalloprotease encoding genes in the *Bsal* genome assembly v2.0 and their matching genes and classifications according to *Farrer et al*. 2017 (R. A. Farrer et al., 2017). Contig numbers (terminal numbers of scaffolds), gene IDs, gene IDs in *Farrer et al*. 2017, IDs of upstream flanking genes and repeats (upstream ID), IDs of downstream flanking repeats and genes (downstream ID), classification as secreted (secreted) or not (NA), Tribes of *Bsal* assembly v2.0, the matching clades in Fig. 5 and M36 clades G1M36 or G2M36 designation (according to *Farrer et al*. 2017) of all M36 metalloproteases in *Bsal* assembly v2.0.

**Table S8.**Gene IDs for genes with a secretion signal, M36 metalloproteases, core-conserved genes (CCGs) and short secreted proteins (SSPs) for *B*.*salamandrivorans* and *B. dendrobatidis*. SSPs IDs are for SSPs defined as being shorter than 200 amino acids and with ≥ 4 cysteines.

## Methods

### Sequencing and library preparation

*Bsal* zoosporangia and zoospores were cultured in tryptone-gelatin hydrolysate-lactose (TgFl) broth in cell culture flasks at 18°C. 200ml of 6 days old cultures were harvested and centrifugated at 1700g for 10 mins at 4°C. Cell pellet was washed with ice cold water and snap frozen in liquid nitrogen. High–molecular weight DNA for Nanopore sequencing was obtained by a customized cetyltrimethylammonium bromide (CTAB) extraction procedure (Schwessinger, 2019; Schwessinger & Rathjen, 2017) with the modification of using RNase A (T3018, NEB) instead of RNase T1. Briefly, cell pellet was ground with a mortar and pestle in liquid nitrogen with 2g of sand, followed by lysis with CTAB, two-step purification with phenol/chloroform/isoamyl alcohol and precipitation with isopropanol. Care was taken to avoid DNA shearing (cut off tips, no heating of samples). DNA concentration was checked using the Qubit BR assay (Invitrogen) and DNA size range profile was checked by Tapestation gDNA screentape (Agilent).

Two independent sequencing libraries were constructed, one with long unfragmented DNA, one with DNA fragmented to 12kb with a gTube (520079, Covaris). DNA ends were FFPE repaired and end-prepped/dA-tailed using the NEBNext FFPE DNA Repair kit (M6630, NEB) and the NEBNext Ultra II End-Repair/dA-tailing Module (E7546, NEB) followed by AMPure XP bead clean-up (A63882, Beckman Coulter). Adapters were ligated using the Genomic DNA by Ligation kit (SQK-LSK109, Oxford Nanopore Technologies) and NEBNext Quick T4 DNA Ligase (E6056, NEB) followed by AMPure XP bead clean-up. The two libraries were successively loaded onto a single PromethION (FLO-PRO002, type R9.4.1) flowcell. The unfragmented library was loaded first. Guppy Basecalling Software v. 3.2.8+bd67289 was used for base calling. A total of 24,402,905 reads were base called and of these 18,678,675 (76.5%) passed the quality check. Passed reads contained 63.78 Gb of DNA sequence (85% of the total DNA nucleotide bases sequenced) amounting to ∼868X depth of coverage. The mean length of nanopore read was 3,415 nt, with an N_50_ of 9,248 and a Mean Read Quality of 10.2. The longest read was 318,012nt long.

### Genome assembly and quality control

Nanopore reads were trimmed using PoreChop v.0.2.3_seqan2.1 (Wick et al., 2017) with default parameters, and filtered where < 500 bp or average read quality > 10 using NanoFilt v.2.6.0 (De Coster et al., 2018). Canu v.1.8 (Koren et al., 2017) was used to assemble reads ≥ 100 kb (∼13X coverage) with stopOnLowCoverage=0.5, genomeSize=0.6g and minReadLength=500 (assembly name = V2 Canu default settings) or with additional parameters corMhapFilterThreshold=0.0000000002 corMhapOptions=“-- threshold 0.80 --num-hashes 512 --num-min-matches 3 --ordered-sketch-size 1000 -- ordered-kmer-size 14 -- min-olap-length 2000 --repeat-idf-scale 50” mhapMemory=60g mhapBlockSize=500 ovlMerDistinct=0.975 (assembly name = V2 Canu non-default settings). Raven v1.1.10 (Vaser & Sikic, 2019) was used to assembly all reads ≥50 kb (∼83X coverage) with default parameters (assembly name = V2 Raven default settings). Medaka v.1.0.3 (https://github.com/nanoporetech/medaka; default parameters) and the trimmed nanopore reads were used for polishing. The polished assembly (V2 Canu default settings Medaka polished) and the unpolished assembly (V2 Canu default settings) were filtered for contigs ≤500bp and corrected with Illumina paired-end sequence data (R. A. Farrer et al., 2017) using Pilon v1.2 (Walker et al., 2014).

We compared the previously published assembly for *Bsal* (assembly name = V1) (R. A. Farrer et al., 2017) to our new assemblies using a variety of tools and metrics (**Table S1**). Assembly quality was assessed using Quast v.5.0.2 (Mikheenko et al., 2018). We evaluated each assembly for pre-annotation gene completeness using Tblastn (-e 1e-10 -v 5 -b 5 -F F) to the 248 Core Eukaryotic Genes (CEG) (Parra et al., 2007) and BUSCO v4.1.1 (Seppey et al., 2019) analysis (datasets eukaryote_odb10 and fungi_odb10). Reapr v1.0.18 (Hunt et al., 2013) was used on the assemblies with Illumina paired-end sequence data (insert size: 441). Internal duplication was assessed by MUMMER v4.0.0beta2 (Kurtz et al., 2004) nucmer (parameters --coords --maxmatch –nosimplify). Non-self-hits (>500bp and >99% identity) were flagged as possible duplications. Dnadiff was run for comparative quantifications of duplications and gaps identified by MUMMER. While all V2 *Bsal* genome assemblies were improvements in multiple metrics compared with V1, we chose the V2 Canu default assembly polished with Pilon only for all subsequent analysis based on high accuracy, contiguity, completeness and coverage (**Supplemental Material**).

### Gene annotation

Gene annotation on the repeat masked V2 assembly was guided by our previous 14.4Gb *Bsal in vitro* RNAseq (NCBI BioProject PRJNA326249) using the Braker2 (Hoff et al., 2019) pipeline (parameters --fungus, --softmasking), which uses Samtools v.0.1.19-44428cd (Li et al., 2009), Bamtools v.2.4.0 (Barnett et al., 2011), Diamond v.2.0.4 (Buchfink et al., 2014), Genemark-ET v4.15 (Lomsadze et al., 2014), and Augustus v.3.2.1 (Stanke et al., 2008). The pipeline identified 11,929 genes, from which 92.74% core eukaryotic genes could be identified via BLASTP (e-value < 1e^-10^). Next, using the Broad Institute’s Vesper annotation pipeline, the genome was BLASTx against Swiss-Prot (Bairoch & Apweiler, 2000) and KEGG (Kanehisa & Goto, 2000), and HMMER hmmscan (Finn et al., 2011) against PFAM (Finn et al., 2014). We ran tRNAscan (Lowe & Eddy, 1997) and RNAmmer (Lagesen et al., 2007) to identify non-protein-coding genes. M36 genes from the V1 assembly were blasted to the Braker2 softmasked predictions, and included in our gene set.

Gene predictions were checked for a variety of issues, including overlapping of noncoding genes, overlapping of coding genes, and the presence of in-frame stops. Genes were named according to evidence from BLASTx and HMMER in the following order of precedence: (i) Swiss-Pro t(Bairoch & Apweiler, 2000), (ii) TIGRfam (Haft et al., 2003), and (iii) KEGG (Kanehisa & Goto, 2000) (where BLASTx hits must meet the 70% identity and 70% overlap criteria to be considered a good hit and for the name to be applied). Otherwise, genes were classified as hypothetical proteins. Genes were functionally annotated by assigning PFAM (release 27) domains (Finn et al., 2014), and BLASTx for KEGG assignment (each where E-value <1×10^−10^), as well as ortholog mapping to genes of known function. SignalP 4.0 (Petersen et al., 2011) and TMHMM (Krogh et al., 2001) were used to identify secreted proteins and transmembrane proteins, respectively.

The protease composition of each chytrid was determined using top high scoring pairs from BLASTp searches (e-value < 1e^-5^) made to the file ‘pepunit.lib’, which is a non-redundant library of protein sequences of all the peptidases and peptidase inhibitors that are included in the MEROPS database (Release 12.1), and compared to the 447 thousand protein sequences in the 2014 version we used in our previous genomic analyses (R. A. Farrer et al., 2017). All proteases with matches to M36 metalloproteases were aligned using MUSCLE v3.8.31 (Edgar, 2004) and trimmed of excess gaps using trimAl 1.2rev59 (Capella-Gutiérrez et al., 2009) gappyout. We constructed the gene trees with RAxML v7.7.8(Stamatakis, 2006) using the JTT amino acid transition model, which was visualized using iTOL v6 (Letunic & Bork, 2021).

Secreted proteins were predicted in each of the 22 chytrid species using SignalP 4.0 (Petersen et al., 2011). These gene sequences had their secretion signal cleaved according to the predicted cleavage site, which were then clustered according to sequence similarity using MCL (http://micans.org/mcl/man/clmprotocols.html) with recommended settings ‘-I 1.4’. Secreted families were classified using PFAM domains (release 34.0) (Finn et al., 2014). Small secreted proteins (SSPs) were classified as those secreted proteins with <300 amino acids and >4 cysteines.

### Chytridiomycota genomic and phylogenetic analysis

The genomes and gene annotation for *B. dendrobatidis* (Bd) JEL423, *S. punctatus* (Sp) and *H. polyrhiza* (Hp) (R. A. Farrer et al., 2017) were downloaded from NCBI (BioProject PRJNA13653, PRJNA37881 and GenBank AFSM00000000 respectively) and FigShare (R. Farrer, 2016). Nineteen further chytrid genomes were downloaded from the Mycocosm portal of the US Department of Energy (DOE) Joint Genome Institute (JGI) (Grigoriev et al., 2014) including *B. helicus* (Ahrendt et al., 2018), *C. hyalinus* JEL632, *C. lagenaria* Arg66, *Chytriomyces sp. nov*. MP71, *E. helioformis* JEL805, *F. jonesii* JEL569, *G. haynaldii* MP57, *G. pollinis-pini* Arg68, *G. prolifer a*(Chang et al., 2015), *G. semiglobifer* Barr 43, *G. variabilis* JEL559, *H. curvatum* SAG235-1, *O. mucronatum* JEL802, *P. hirtus* BR81, *R. globosum* JEL800 (Mondo et al., 2017), *T. arcticum* BR59, *C. replicatum* JEL714 (Mozley-Standridge et al., 2009) and *C. polystomum* WB228. We excluded *B. helicus* from further analysis as it had <75% complete BUSCOs, suggesting the assembly or gene calls are incomplete.

Single copy orthologs were identified between chytrids using the Synima (R. A. Farrer, 2017) pipeline with Orthofinder, and aligned using MUSCLE v3.8.31 (Edgar, 2004). A maximum likelihood tree was constructed using IQ-Tree v1.6.12 (Nguyen et al., 2015, p.) with the LG amino acid substitution model (the best fitting model according to ProtTest v3.4.2 (Darriba et al., 2011)) with 1000 ultrafast bootstraps, and visualized using Figtree v1.4.4 with midpoint rooting.

### Repeat Analysis

Repeat content was identified using Repeatmodeller v.2.0.1 (Flynn et al., 2020) with rmblast v.2.10.0+ and Tandem Repeat Finder v.4.09(Benson, 1999), RepeatScout v.1.06 and RepeatMasker v.4.0.5 (Smit et al., 2015). The output of Repeatmodeller (consensi.fa.classified) was then used as a library for RepeatMasker. The repeat content and family distribution for each chytrid species was determined from RepeatMasker.out, excluding lower scoring matches whose domain partly (<80%) includes the domain of another match.

TE and repeat distributions in the genome were assessed using Repeatmasker .gff outfiles and visualized using IGV 2.8.2 (Robinson et al., 2011). TE distribution in relation to GC content was analyzed using Pilon’s GC.wig files. Additionally, TE and repeat content of 10kb windows assigned to different quadrants was calculated using custom scripts based on RepeatMasker .out files & lists of genes assigned to quadrants.

To assess if repeat content was correlated to genome assembly quality, Spearman’s Rank Correlation Coefficients, Spearman’s correlation and linear regression (linear model fitting based on formula by Wilkinson and Rogers (1973) (Wilkinson & Rogers, 1973)) were calculated between repeat content and N_50_, the number of contigs, genome length, as well as the number of genes using ggpubr (https://github.com/kassambara/ggpubr).

For the heatmap showing repeat superfamily profiles, heatmap.2 was used with hierarchical clustering and Euclidian as a distance measure. Repeat families that did not exceed 1% abundance in any of the chytrids were excluded. Repeat families in *Bd* and *Bsal* were aligned using blastn with an e-value of 0.01, no filtering (-dust no, -soft_masking false) and a wordsize of 7 to determine homologs. Telomeric sequences were manually curated.

### Genome compartmentalization analysis

Flanking intergenic distance was calculated for all non-terminal protein coding genes. For each chytrid species, the median distance was used define four quadrants: Density plots of intergenic distances were constructed for all non-terminal protein coding genes, and several gene subsets including genes with a secretion signal, SSPs, conserved chytrid BUSCO genes and M36 metalloproteases. The median 5’ and 3’ intergenic distance for all protein coding genes in a given species was used to define four quadrants including bottom left (gene-rich/repeat-sparse), top right (gene-poor/repeat-rich), bottom right (long 3’ intergenic distance, short 5’ intergenic distance) and top left (short 3’ intergenic distance, long 5’ intergenic distance).

To identify enrichment of gene categories in each quadrant, Hypergeometric tests were performed on all genes, the aforementioned four gene categories, and the largest 10 secreted families (determined by MCL). Hypergeometric tests were also used to determine enrichment for flanking repeat families and to determine whether genes falling in one of the quadrants are enriched on certain chromosomes compared to the overall distribution of genes in quadrants on all chromosomes. Critical *p*-values for hypergeometric and Х^2^ enrichments were determined using Bonferroni correction with an α-level of 0.01. For gene category enrichment, we performed 16 tests = 0.00063. For secreted families and quadrant enrichments tests on chromosomes, we performed 4 tests = 0.0025. For flanking repeat families, the correction was adjusted based on the total number of repeat families flanking.

There is currently no known population structure of *Bsal*, thereby precluding the study of intra-population genetic variation. However, there are multiple lineages of *Bd* described (R. A. Farrer et al., 2011; O’Hanlon et al., 2018) and therefore genetic variation for this species can be compared to intergenic distance. Paired-end Illumina data from representatives of all five known lineages (*Bd*GPL JEL423, *Bd*CAPE TF5a1, *Bd*CH ACON, *Bd*Asia-1 KRBOOR_323, *Bd*Asia-2 CLFT065, and a hybrid of unknown parentage SA-EC3) were obtained from the NCBI Sequence Read Archive (SRA) (R. A. Farrer et al., 2011, 2013; O’Hanlon et al., 2018). The Genome Analysis Toolkit (GATK) v.4.1.2.0 (McKenna et al., 2010) was used to call variants. Our Workflow Description Language (WDL) scripts were executed by Cromwell workflow execution engine v.48 (Voss et al., 2017). Briefly, raw sequences were pre-processed by mapping reads to the reference genome *Bd* JEL423 using BWA-MEM v.0.7.17 (Li, 2013). Next, duplicates were marked, and the resulting file was sorted by coordinate order. Intervals were created using a custom bash script allowing parallel analysis of large batches of genomics data. Using the scatter-gather approach, HaplotypeCaller was executed in GVCF mode with the diploid ploidy flag. Variants were imported to GATK 4 GenomicsDB and hard filtered (QD < 2.0, FS > 60.0, MQ < 40.0, GQ ≥ 50, AD ≥ 0.8, DP ≥ 10). The direction and magnitude of natural selection for each lineage was assessed by measuring the rates of non-synonymous substitution (*dN*), synonymous substitution (*dS*) and omega (ω = *dN*/*dS*) using the yn00 program of PAML (Yang, 2007) implementing the Yang and Nielsen method, taking into account codon bias (Yang & Nielsen, 2000). The program was run on every gene in each isolate using the standard nuclear code translation table. Hypergeometic tests were calculated for genes with ω > 1 in each quadrant. We performed 20 tests per lineage, thus the *p*-value was Bonferroni corrected to 0.0005 at an α-level of 0.01

Х^2^ enrichments tests of independence were performed on a range of genes including 1) genes with a secretion signal, 2) SSPs, 3) conserved chytrid BUSCO genes, 4) repeat families flanking genes coding for secreted proteins, 5) M36 metalloproteases and 6) *Bd* genes that have ω > 1. Briefly, 2×2 contingency tables were generated for each test, comprising two groups of dichotomous variables (number of genes in or not in the gene category of interest, and the number of genes in or not in a given quadrant). X^2^-tests for goodness of fit were performed to determine whether the distribution of genes within each quadrant was significantly different from the expected distribution (25% each). X^2^-tests were performed on each contig iteratively to test for genomic hot-spots for rapid evolution.

Wilcoxon rank-sum tests were computed to test the null hypothesis that the log_10_ mean intergenic distances of the feature category of interest (SSPs, HKGs, M36 metalloproteases and secreted proteins only) and the log_10_ mean intergenic distances of all the other genes that are either a) not in that feature category of interest or b) of a different feature category have the same continuous distribution. Wilcoxon rank-sum tests were performed using Rstatix v0.7 (https://cran.r-project.org/web/packages/rstatix/index.html) wilcox_test (conf.level=0.95). Wilcoxon effectsize was determined using Rstatix wilcox_effsize (conf.level=0.95,nboot=1000 and ci=TRUE). For Wilcoxon rank sum tests, the adjusted p-values in the violin plots for α-levels of 0.01 to 0.0001 and 6 tests were as follows: ns: p > 0.0017, *: p <= 0.0017, **: p ≤ 0.00017, ***: p ≤ 1.7E-5, ****: p ≤ 1.7E-6.

Consecutive gene counts were generated using lists of genes assigned to their quadrants as defined above and a bespoke script. To assess the significance of finding a certain number k of consecutive genes of the same quadrant, discrete pattern markov chains were used. The probability of transitioning from one quadrant to the next was set to 0.25. Based on that, a (k+1) X (k+1) transition matrix was generated. Once the transition matrix was constructed, for a given value of *n* the probability of having the number of consecutive genes of a certain quadrant in the chain was *P*(*W*|*n*) = {*Pn*}0,_k_. In the calculation, n was set to 100 repetitions of equiprobable outcome. *W* is the event of the occurrence of k consecutive genes of the same quadrant.

## Data availability

Raw *B. salamandrivorans* sequences are deposited at GenBank under Bioproject PRJNA666901. The genome assembly and annotations for *Bsal* are deposited at GenBank under Bioproject PRJNA666901.

## Acknowledgments

RAF and NH are supported by MRC grant MR/N006364/2 and a Wellcome Trust Seed Award (215239/Z/19/Z). TW is supported by an MRC Studentship (112189-G). This project utilised equipment funded by the UK Medical Research Council (MRC) Clinical Research Infrastructure Initiative (award number MR/M008924/1).

## Notes

### Competing Interest Statement

The authors have declared no competing interest.

