## Supplementary figures and images for "Two-speed genome expansion drives the evolution of pathogenicity in animal fungal pathogens"

### Supplemental Figure 1

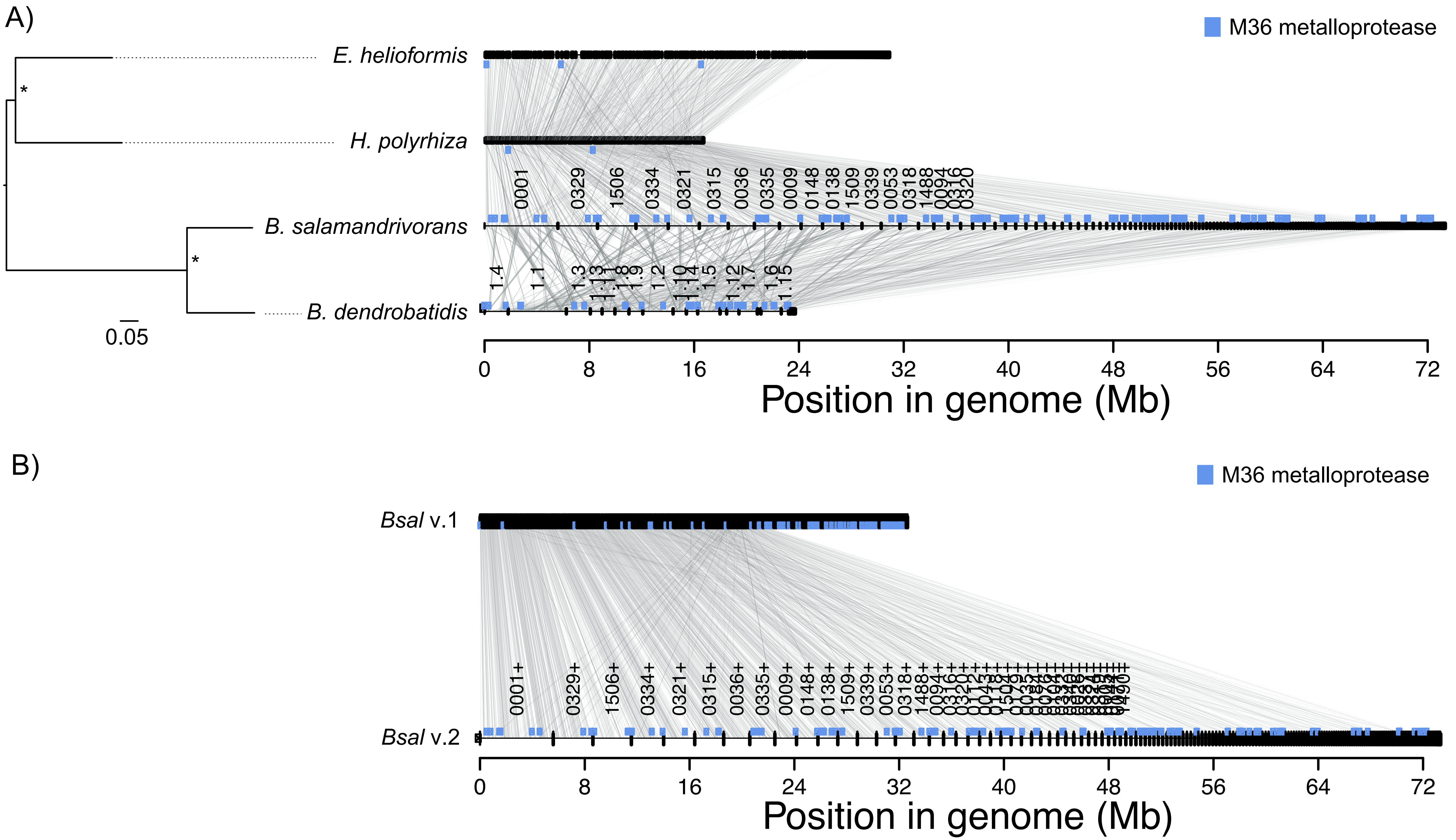

### Supplemental Figure 2

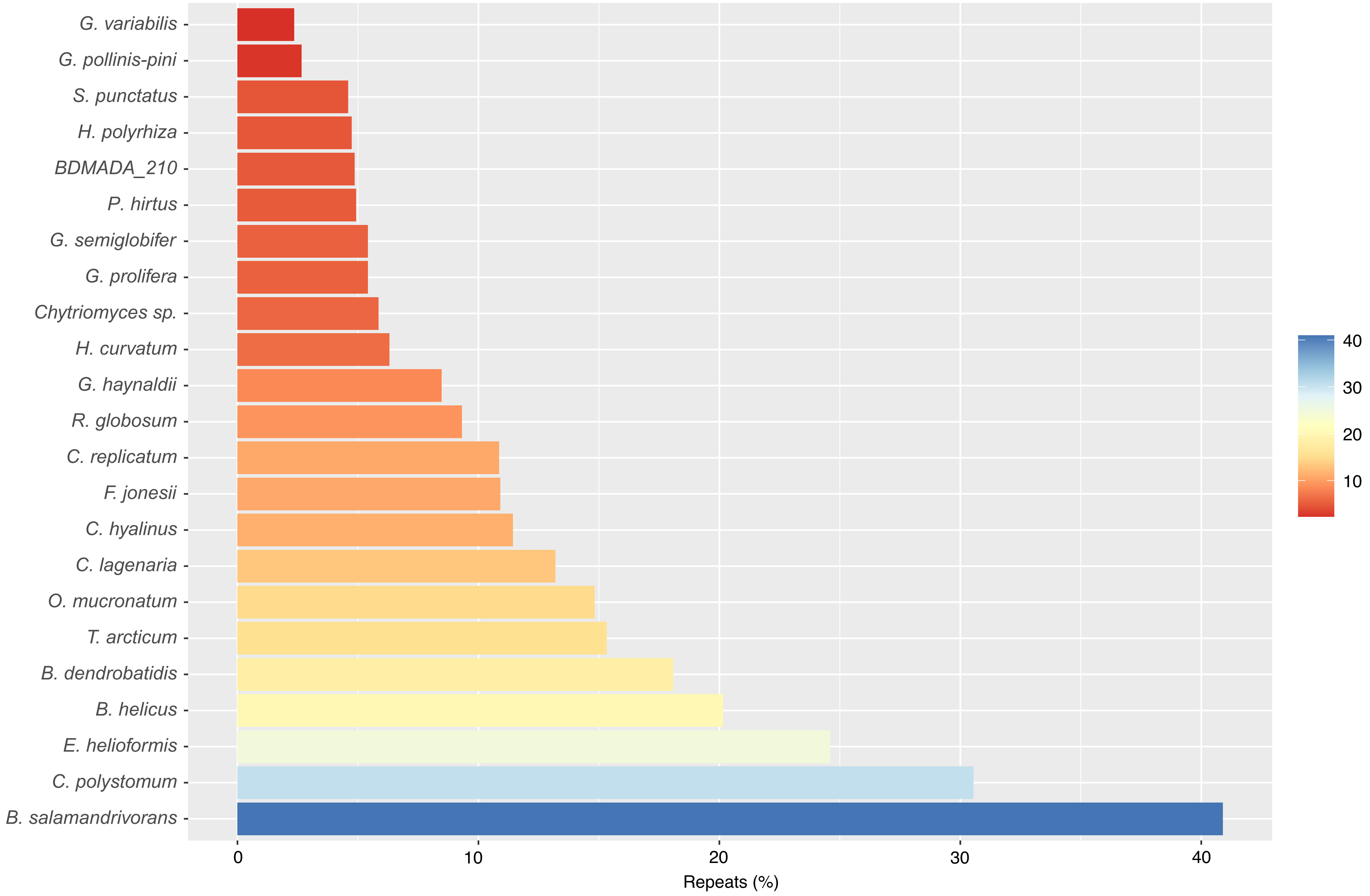

### Supplemental Figure 3

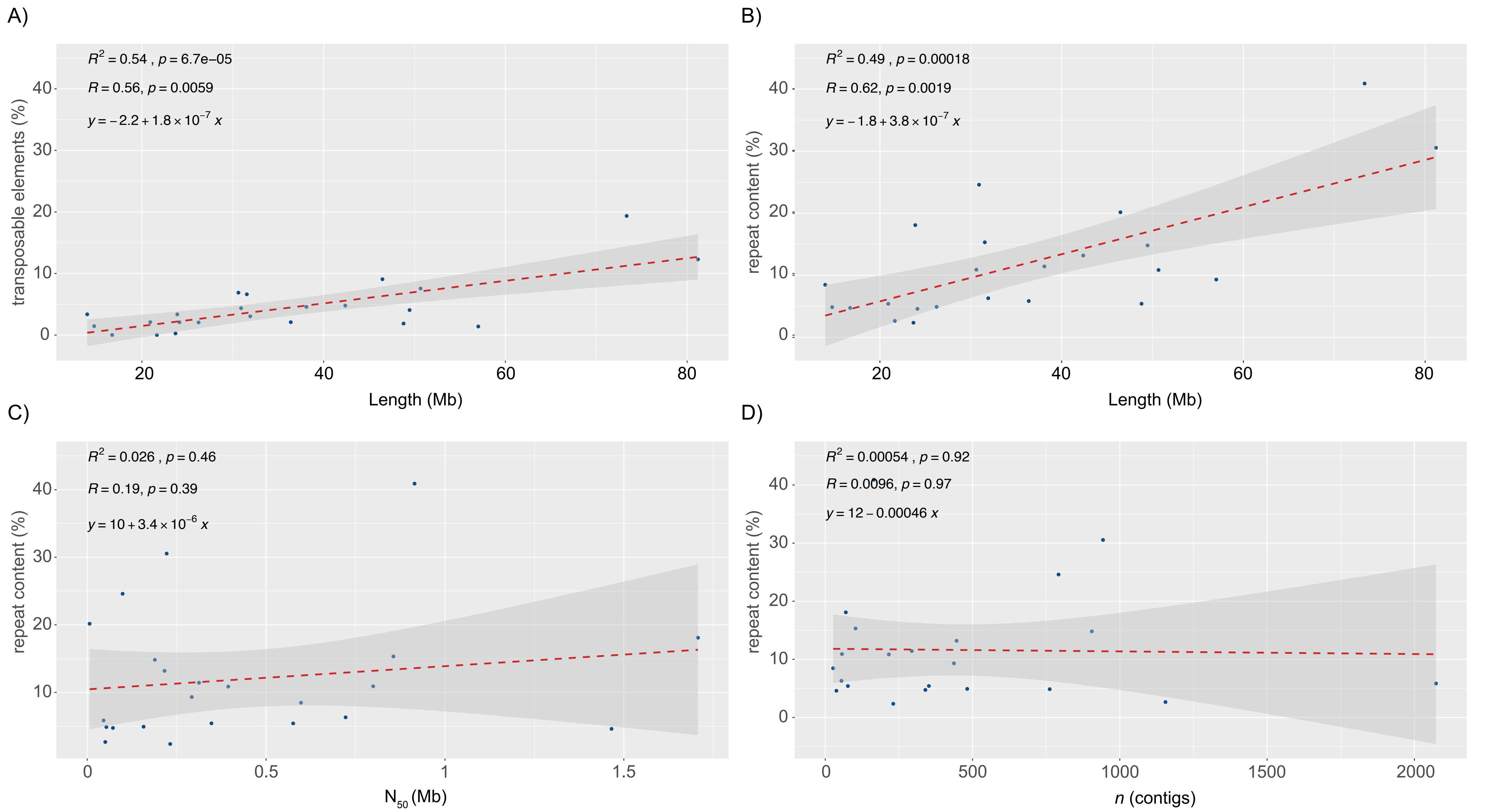

### Supplemental Figure 4

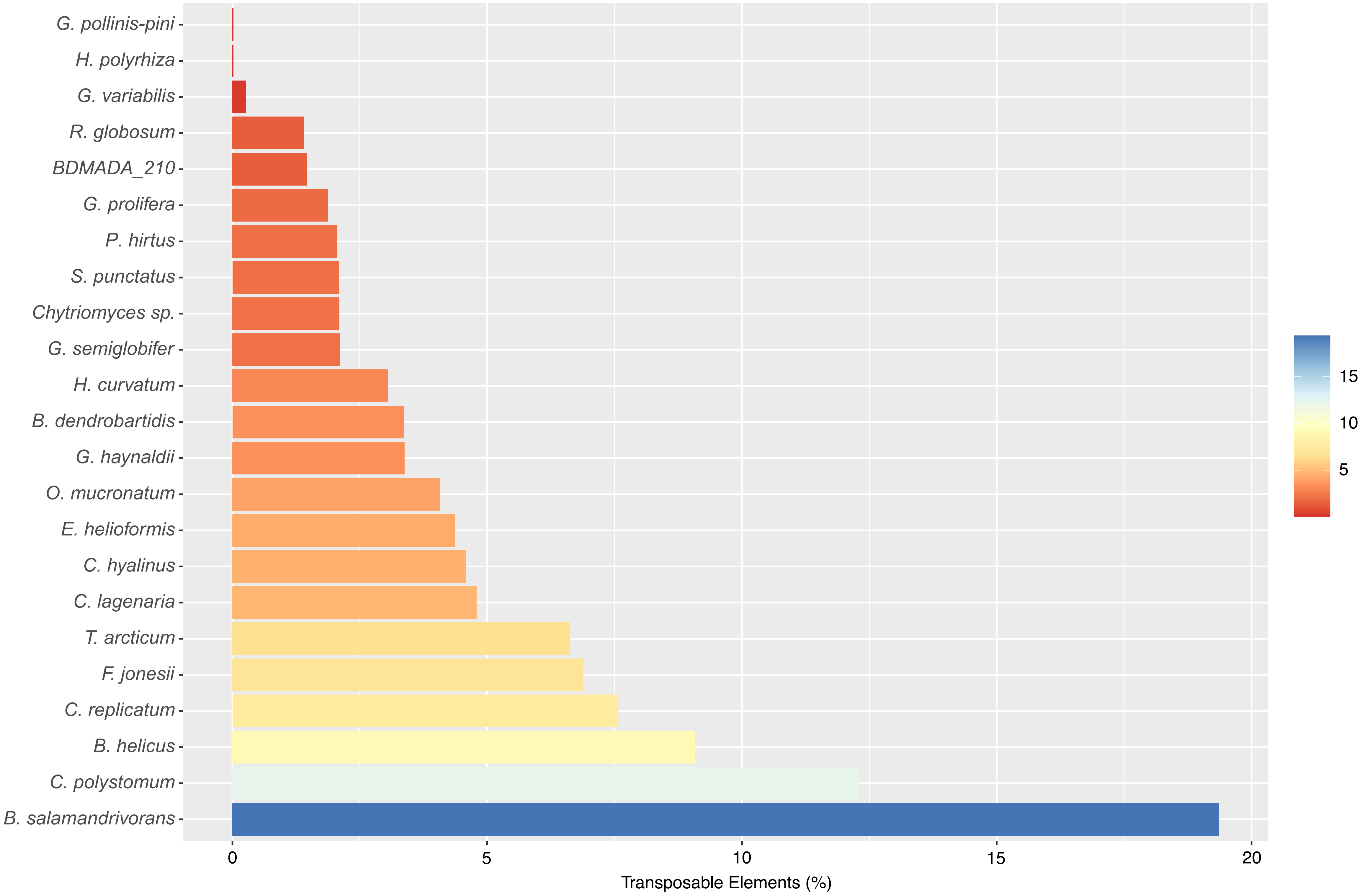

### Supplemental Figure 5

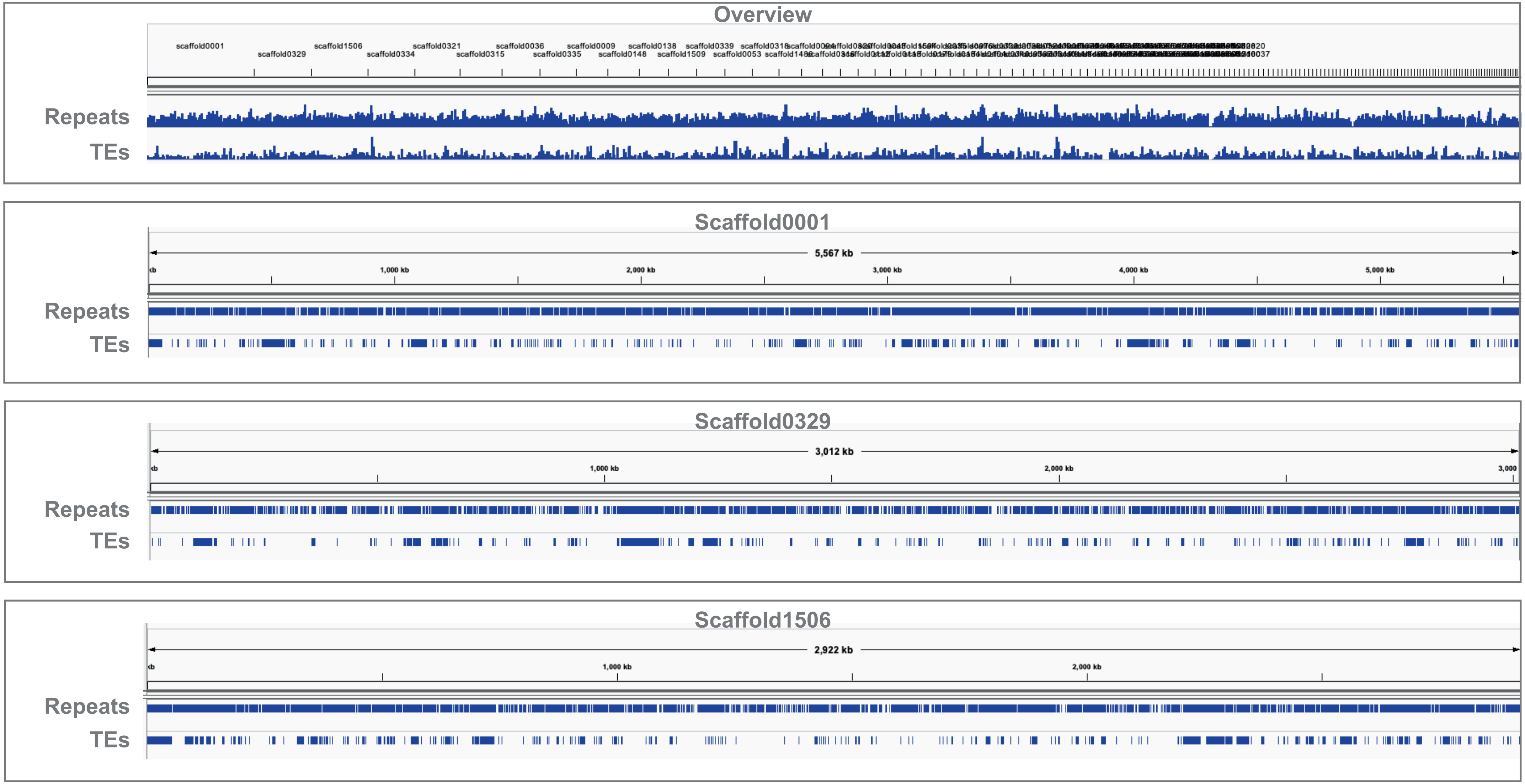

### Supplemental Figure 6

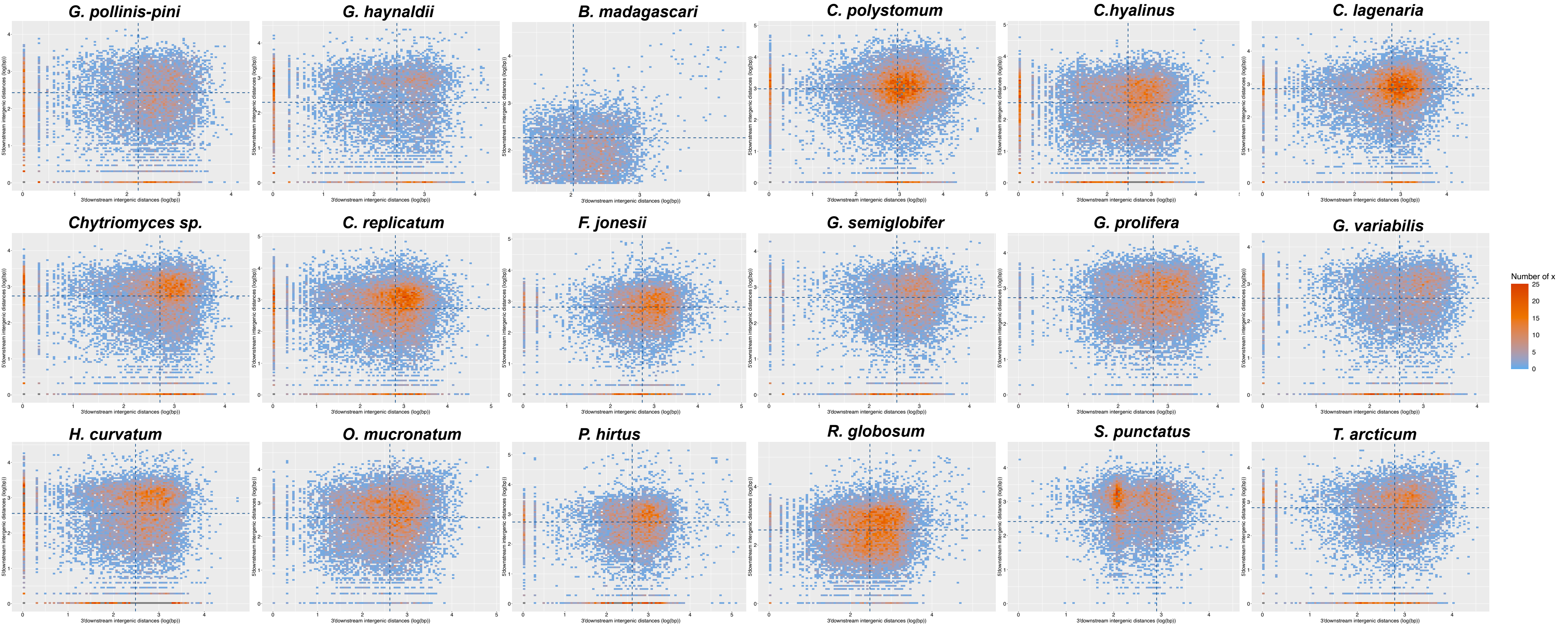
